# Data-driven Assessment of Structural Image Quality

**DOI:** 10.1101/125161

**Authors:** Adon F. G. Rosen, David R. Roalf, Kosha Ruparel, Jason Blake, Kevin Seelaus, Lakshmi P. Villa, Rastko Ciric, Philip A. Cook, Christos Davatzikos, Mark A. Elliott, Angel Garcia de La Garza, Efstathios D. Gennatas, Megan Quarmley, J. Eric Schmitt, Russell T. Shinohara, M. Dylan Tisdall, R. Cameron Craddock, Raquel E. Gur, Ruben C. Gur, Theodore D. Satterthwaite

## Abstract

Data quality is increasingly recognized as one of the most important confounding factors in brain imaging research. It is particularly important for studies of brain development, where age is systematically related to in-scanner motion and data quality. Prior work has demonstrated that in-scanner head motion biases estimates of structural neuroimaging measures. However, objective measures of data quality are not available for most structural brain images. Here we sought to identify quantitative measures of data quality for T1-weighted volumes, describe how such measures of quality relate to cortical thickness, and delineate how this in turn may bias inference regarding associations with age in youth. Three highly-trained raters provided manual ratings of 1,840 raw T1-weighted volumes. These images included a training set of 1,065 images from Philadelphia Neurodevelopmental Cohort (PNC), a test set of 533 images from the PNC, as well as an external test set of 242 adults acquired on a different scanner. Manual ratings were compared to automated quality measures provided by the Preprocessed Connectomes Project’s Quality Assurance Protocol (QAP), as well as FreeSurfer’s Euler number, which summarizes the topological complexity of the reconstructed cortical surface. Results revealed that the Euler number was consistently correlated with manual ratings across samples. Furthermore, the Euler number could be used to identify images scored “unusable” by human raters with a high degree of accuracy (AUC: 0.98-0.99), and outperformed proxy measures from functional timeseries acquired in the same scanning session. The Euler number also was significantly related to cortical thickness in a regionally heterogeneous pattern that was consistent across datasets and replicated prior results. Finally, data quality both inflated and obscured associations with age during adolescence. Taken together, these results indicate that reliable measures of data quality can be automatically derived from T1-weighted volumes, and that failing to control for data quality can systematically bias the results of studies of brain maturation.

## INTRODUCTION

In-scanner motion and other artifacts are increasingly appreciated as a source of bias in neuroimaging research. In-scanner motion reduces image quality, and is also related to subject characteristics of interest, including participant age (Power et al., 2012; Satterthwaite et al., 2012). As such, it has the potential to systematically confound inference, especially in studies of lifespan development (Zuo et al., 2017). While motion has long been a well-described methodological obstacle in medical imaging (Bellon et al., 1986; Smith and Nayak, 2010), and a known confound for task-related fMRI (Friston et al., 1996), it has recently attracted additional scrutiny. Following reports that even small amounts of in-scanner motion can bias studies of functional connectivity (Power et al., 2012; Satterthwaite et al., 2012; Van Dijk et al., 2012), there has been a proliferation of recent studies that have documented the impact of data quality on other imaging modalities, including T1-weighted neuroimaging of brain structure (Alexander-Bloch et al., 2016; Pardoe et al., 2016; Reuter et al., 2015; Savalia et al., 2017).

Following initial work to assess motion’s impact on structural images (Atkinson et al., 1997), much subsequent work has addressed structural image quality issues driven by scanner and platform-related variation (Chen et al., 2014; Magnotta and Friedman, 2006; Styner et al., 2002; Woodard and Carley-Spencer, 2006). However, several published studies have used unique attributes of T1-weighted images to quantify image quality. Specifically, Mortamet et al. (2009) introduced a quality index (Qi) that accurately identified unusable volumes (AUC=0.93) collected as part of the Alzheimer’s Disease Neuroimaging Initiative. Furthermore, Pizarro et al. (2016) developed statistics based on specific artifacts such as eye motion, ringing and tissue contrast. Combined in a multivariate approach, these statistics classified unusable volumes with a classification accuracy of 80%. However, these studies examined neither quality indices related to measures of brain structure, nor how quantitative indices of data quality might be used to account for biases in group level analyses. This is particularly relevant given that measures of brain structure such as cortical thickness are frequently used as putative biomarkers in research on development, aging, and a myriad of neuropsychiatric diseases.

Research using functional timeseries has typically summarized motion via the “framewise displacement” calculated from timeseries realignment parameters (Power et al., 2012; Satterthwaite et al., 2012; Van Dijk et al., 2012). However, most structural imaging sequences do not provide a ready estimate of participant motion during acquisition. A variety of motion-tracking systems have recently become widely available for use in structural MRI, including in-bore optical systems as well as approaches using the MRI scanner itself to track motion, allowing for motion to be directly quantified in a manner akin to functional imaging time series (Zaitsev et al., 2015). Reuter et al. (2015) used the vNav-MPRAGE sequence (Tisdall et al., 2012), which simultaneously acquires a T1-weighted volume and performs motion tracking with the MRI scanner, to demonstrate in 12 healthy adults that motion during the T1 sequence was associated with spurious alterations of cortical thickness and cortical volume. Tisdall et al. (2016) demonstrated that using this motion information prospectively could substantially reduce the deleterious effects of motion on both image quality and subsequent morphometry.

Despite the clear importance of such work, the vast majority of T1-weighted imaging sequences acquired to date lack any motion-tracking or motion-correction technology, and thus cannot derive a quantitative assessment of motion. While current commonly-used processing pipelines (including CCS, DPABI, and HCP pipelines, Marcus et al., 2013; Xu et al., 2015; Yan et al., 2016) provide a range of measures of data quality for functional timeseries, validated quantitative measures of data quality are not typically produced for the T1 volume. Accordingly, three important recent studies used motion during a functional imaging sequence acquired during the same scanning session as a proxy of in-scanner motion during the structural scan (Alexander-Bloch et al., 2016; Pardoe et al., 2016; Savalia et al., 2017). This approach is based on the observation that participant motion tends to be highly correlated across acquisitions: individuals with high motion in one sequence tend to have high motion in other sequences (Pardoe et al., 2016; Yan et al., 2013). Three studies demonstrated that higher motion during a functional sequence acquired in the same session is associated with cortical thickness, even in those scans which passed manual quality assurance procedures (Alexander-Bloch et al., 2016; Pardoe et al., 2016; Savalia et al., 2017). Furthermore, Salvia et al. (2017) demonstrated that unaccounted-for motion artifact inflated the apparent effects of aging. While motion during a functional sequence is an opportune proxy for motion during a structural scan, it nonetheless has several limitations. First, it requires that a functional scan was acquired, which may not be possible due to subject factors, time restrictions, or study design. Second, the ecological validity of the proxy is likely to vary with ordering effects, amount of time between scans, as well as other uncontrolled variables such as patient comfort.

In this study, we sought to identify quantitative measures of data quality that could be derived from the T1 volume alone. Measures of data quality were provided by the Preprocessed Connectomes Project’s Quality Assurance Protocol (QAP); the Euler number provided by FreeSurfer was also evaluated. We investigated the degree to which these quantitative measures could be used to identify unusable images, and compared them to proxy measures of data quality provided by functional sequences. Furthermore, we described how quantitative metrics of image quality related to cortical thickness, and potentially confound associations with age. Throughout, we leveraged the large sample provided by the Philadelphia Neurodevelopmental Cohort (PNC), as well as an independent sample of adults imaged on a different scanner. As described below, we found that measures derived from the T1-weighted volume provide useful measures of image quality.

## METHODS

### Approach overview

Our overall goal was to evaluate quantitative measures of image quality directly from structural MRI volumes. This process included several discrete tasks. First, all image analysts underwent rigorous training, and then independently rated all images. Second, we evaluated quantitative measures of image quality to determine which aligned best with manual ratings. Third, we used these quantitative measures to identify images that were unusable; we refer to this as the “inclusion” model. Fourth, we compared this approach to proxy measures estimated from motion during functional time series acquired during the same session. Fifth, we examined how quantitative measures of image quality related to cortical thickness as measured by the popular FreeSurfer platform (Fischl and Dale, 2000). Sixth and finally, we examined how data quality might bias inference regarding associations with age in samples of youth. All analysis code is publicly available here <u>https://github.com/PennBBL/RosenT1QA</u>.

### Participants

We included a total of 1,840 images across two studies that used different scanners **(Table 1**). This included 1,598 images from the PNC (Satterthwaite et al., 2014) as well as an additional 242 images from a study acquired on a different scanner (Roalf et al., 2015). Specifically, 1,065 PNC images were used for training, and 533 were used during testing. In order to maintain a similar distribution of age, sex, and manual image quality rating across the training and testing samples of the PNC, we used the ‘caret’ package in R (Kuhn et al., 2016). The data from the second study were used only as an external test dataset. This second cohort was comprised of adults, and thus not matched on demographic details (see Table 1).

**Table 1:** Demographic information of the training and validation datasets.

| Study | N | % Female | Age Mean | Age SD |
| --- | --- | --- | --- | --- |
| Training | 1065 | 51 | 14.90 | 3.70 |
| Testing 1 | 533 | 44 | 15.10 | 3.68 |
| Testing 2 | 242 | 48 | 41.36 | 16.99 |

### Image acquisition

All imaging data from the PNC were acquired on the same 3T Tim Trio scanner with a 32-channel head coil (Siemens: Erlangen, Germany) as previously described (Satterthwaite et al., 2014). Structural images were acquired using a magnetization-prepared, rapid-acquisition gradient-echo (MPRAGE) T1-weighted sequence (TR = 1810ms; TE = 3.51ms; T1 = 1100ms; FoV = 180 × 240mm; flip angle = 9°; GRAPPA factor = 2; BW/pixel = 130 Hz; resolution: 0.94mm x 0.94mm x 1.0mm; Acquisition time = 3:28). Prior to scanning, in order to acclimate participants to the MRI environment and to help subjects learn to remain still during the actual scanning session, a mock scanning session was conducted using a decommissioned MRI scanner and head coil. Mock scanning was accompanied by acoustic recordings of the noise produced by gradient coils for each scanning pulse sequence. In the external test set, T1-weighted volumes were collected on a different 3T Tim Trio scanner, using an 8-channel head coil with the following acquisition parameters: TR = 1680ms; TE = 4.67ms; T1 = 1100ms; FoV = 180 × 240mm; flip angle = 15°; bandwith/pixel = 150Hz; resolution: 0.94mm x 0.94mm x 1.0mm; acquisition time = 5:00 (Roalf et al., 2015).

### Image Processing

Cortical reconstruction of the T1 image was performed for all subjects using FreeSurfer version 5.3 (Fischl, 2012). FreeSurfer includes registration to a template, intensity normalization, gray and white matter segmentation, and tessellation of the gray/CSF and white/gray matter boundaries (Dale et al., 1999); cortical surfaces are inflated and normalized to a template via a spherical registration. Cortical thickness is measured as the shortest distance between the pial and the white matter tessellated surfaces (Dale et al., 1999). The cortex was then parcellated into 40 regions (Desikan e al., 2006) and cortical thickness was averaged across parcels to obtain regional cortical thickness estimates.

### Manual rating procedure and rater training

Similar to prior work (Reuter et al., 2015; Savalia et al., 2017), all images were rated on quality using a 0-2 ordinal scale. Initial pilot testing indicated that using systems with more quality classes (i.e., 4 or 5 rating classes) resulted in substantially diminished inter-rater reliability even among experts. In the 3-class framework used, a “0” denoted images that suffer from gross artifacts and were considered unusable. In contrast, a “2” was assigned to images free from visible artifact. The intermediate “1” category was used for images with some artifact, but which still would be considered usable.

A rigorous process of training was used to ensure high inter-rater reliability (see Figure 1). First, anchors and exemplars for the three quality classes were agreed upon through consensus of 5 experts, including a board-certified neuroradiologist (JES), an MR physicist (MAE), a cognitive neuroscientist (DRR), an experienced image analyst (AR), and a neuropsychiatrist (TDS). Next, two of these experts (DRR and TDS) created a larger training sample by rating 100 images independently. Initial concordance was 93%; discrepancies were resolved through consensus, thus yielding a set of 100 images that were used to train three image analysts (KS, PV, JB) who served as the raters for the complete dataset. These three analysts were trained to >85% agreement in this dataset. This required two rounds of blind rating: during the first round, agreement with the expert consensus was 82% (JB), 57% (PV), and 82% (KS). Following further training, each rater re-rated this set of 100 images (presented in a different order, without identifiers), and achieved an accuracy of 91% (JB), 86% (PV), and 94% (KS). Having met reliability benchmarks, these three raters then independently rated all 1,840 images across datasets.

**Figure 1:**
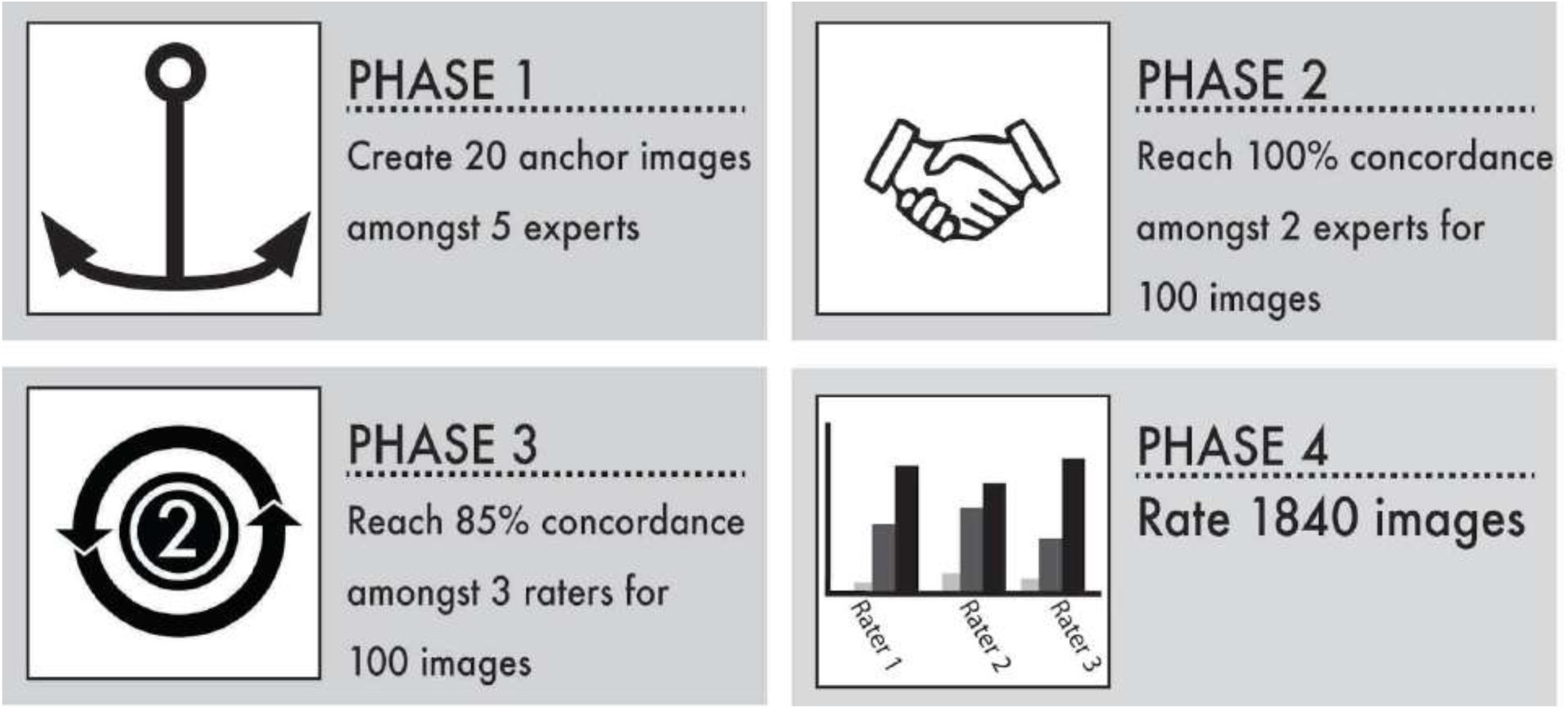
Training protocol for manual raters. There were 4 phases of training. ***Phase 1***: 5 neuroimaging experts reviewed 20 PNC images selected to have various levels of artifact. These images were used to establish rating anchors, which were then used for Phase 2. ***Phase 2***: Two experts (TDS & DRR) rated 100 images. 100% concordance was achieved through consensus. ***Phase 3***: Three new raters were trained on the 100 images used in Phase 2, until the raters achieved 85% concordance after two rounds. ***Phase 4***: All 3 trained raters manually rated 1,840 images across the PNC and the external test dataset (see Table 1).

Rater concordance was evaluated using two measures: the weighted-κ statistic and polychoric correlations. These two measures provide complementary information: while the weighted-κ assesses absolute rating agreement, the polychoric correlation assesses the ordering of the ratings. Variation amongst raters were assessed using a repeated measures ANOVA model. The relationship between manual rating and age was evaluated using partial Spearman’s correlations; sex differences were evaluated using a Wilcoxon signedrank test.

### Quantitative metrics of structural image quality

We evaluated the utility of an array of quantitative imaging measures included in QAP (see Table 2)(Shehzad et al., 2013). QAP version 1.0.3 utilized FMRIB’s Automated Segmentation Tool (FAST, Zhang et al., 2001) for image segmentation, which enables definition and quantification of quality metrics using an image’s gray matter, white matter, and background voxels. Steps were taken to avoid the inclusion of neck and face tissue within the image’s background for the calculation of all background metrics as previously described (Mortamet et al., 2009). In addition to the measures included in QAP, we also calculated image kurtosis and skewness (Joanes and Gill, 1998) for each tissue class and background using tools included in the ‘ANTsR’ (Avants et al., 2016) and ‘psych’ (Revelle, 2017) packages in R; these measures have been integrated into recently-released updates to QAP. Finally, we considered a quality measure produced by the FreeSurfer pipeline: the Euler number (Dale et al., 1999) which is a measure of the topological complexity of the reconstructed cortical surface. Euler number is calculated separately for each hemisphere; we averaged across both hemispheres here to produce one value per subject.

**Table 2:** Quantitative image quality metrics.

| Quantitative Metric | Abbreviation | Citation |
| --- | --- | --- |
| Signal-to-noise ratio | SNR | Magnotta and Friedman, 2006 |
| Contrast-to-noise ratio | CNR | Magnotta and Friedman, 2006 |
| Foreground-to-background energy ratio | FBER | NA |
| Quality index 1 | Qi1 | Mortamet et al., 2009 |
| Image smoothness | FWHM | Friedman et al., 2006 |
| Entropy focus criterion | EFC | Atkinson et al., 1997 |
| Euler number | Euler | Dale et al., 1999 |

In order to visualize the relationship between quantitative measures and manual quality rating, we plotted the mean value for each image quality metric versus the mean manual quality rating. Furthermore, we also calculated partial Spearman’s correlations between the average manual rating and quantitative metrics (while controlling for age, age squared, and sex). For these plots and subsequent analyses, we collapsed any image with an average rating less than 1 into the ‘0’ bin due to the small cell size of these bins.

### Identifying unusable images: the “inclusion” model

A common step in sample construction is to remove images where data quality is so low that the images are considered unusable. We sought to use the quantitative measures of data quality described above to automatically identify unusable images. To do so, we constructed a logistic regression model for each quality metric, where the outcome was a binarized image quality score (i.e., images with a quality score of “0” versus all others). The primary measure of model performance was area under the curve (AUC); accuracy, sensitivity, and specificity were also calculated.

As described below (see Results), a single variable performed quite well in this task. However, in order to ascertain if using additional measures of data quality would aid in classification, we also evaluated multivariate models. Model training began with a simple mass-univariate model and then added features to create a multivariate model in a forward-stepwise manner. The first (base) variable in the multivariate model was defined as the variable with the best performing receiver operator curve (ROC) as measured by area under the curve (AUC) in the mass-univariate analyses conducted in the training sample. Additional measures were added separately to this base model, and the AUC was re-calculated. The best performing feature was selected, and this process was repeated. At each step, in order to determine whether an additional model parameter provided significantly improved classification, we calculated the Delong statistic, which tests for a significant increase in AUC between models (DeLong et al., 1988). Model building was terminated when no significant increase in AUC was found.

After construction of the model in the training set, the classification threshold criterion from the training set was applied to the first (internal) testing dataset as well as the second (external) test set. The same outcome measures (AUC, accuracy, sensitivity, specificity) were then calculated separately for each test sample. Performance using the threshold defined in the training set was compared with outcomes when the classification thresholds were calculated separately for each dataset.

### Comparison to motion in functional scans acquired in the same session

Three recent reports demonstrated that motion in functional sequences acquired during the same scanning session acts as an effective proxy for structural image quality (Alexander-Bloch et al., 2016; Pardoe et al., 2016; Savalia et al., 2017). Accordingly, we next compared our quantitative measure of structural image quality to head motion estimated from functional sequences. This was only conducted in the PNC sample. These additional sequences included a pseudo-continuous arterial spin labeled (PCASL) perfusion scan, two task fMRI scans (tfMRI 1 & tfMRI 2), and one resting functional connectivity scan (rsfMRI) (Satterthwaite et al., 2014, 2016). As sequences acquired at the end of the scanning session are more likely to be missing, we examined motion during each functional sequence, which was summarized as the Frame Displacement (FD), estimated using the average root mean square displacement as calculated by FSL’s MCFLIRT (Jenkinson et al., 2002). Next, we evaluated attrition over the course of the scanning session, and plotted the proportion of missing scans for each sequence, separated by the manual quality rating of the T1 image. Finally, we evaluated the degree to which motion during the functional sequence could identify unusable images using a logistic regression model as described above. In order to ensure that the same sample was considered by each model, this analysis was conducted in a sample of 1,275 PNC subjects that spanned both training and testing samples with complete data across all sequences.

### Relationship of quantitative measures of quality to cortical thickness

As described below, Results revealed that a single metric – the Euler number – was sufficient for identifying unusable images with a high degree of accuracy. Next, we examined associations between this quantitative measure of data quality and cortical thickness in the images that were considered usable (according to their manual rating). Specifically, we used linear regression to examine the association between the Euler number and regional estimates of cortical thickness derived from the FreeSurfer pipeline. In this mass-univariate analysis, cortical thickness was the outcome and Euler number was the predictor of interest; age, age squared, and sex were included in these regression models as covariates. Multiple comparisons across regions were accounted for using the False Discovery Rate (FDR; *q* < 0.05).

### Impact of data quality on associations with age

The analysis described above revealed substantial relationships between data quality and cortical thickness. As a final step, we examined how data quality might bias tests examining associations with age. Accordingly, using the training and testing samples from the PNC, we conducted mediation analyses to determine whether quantitative estimates of data quality (e.g., the Euler number) might mediate the apparent relationship between age and brain structure. Our test statistic for this analysis was the Sobel’s *z*-score (Sobel, 1982), which was calculated for each cortical region. Sobel’s *z*-score estimation was implemented in the ‘bda’ package in R (Wang, 2015). Multiple comparisons were accounted for using FDR as above (*q* < 0.05).

## RESULTS

### Highly trained manual raters achieve good concordance

Across datasets, image quality was relatively high, with a minority of images being considered unusable (Figure 2A-C). Although there were significant differences among raters (training: *F*[2, 3198] = 39.65, *p*<.0001; internal testing: *F*[2, 1599] = 17.74, *p*<.0001; external testing: *F*[2,837] = 3.50, *p*<.05), post-hoc analysis found that raters never disagreed by more than one quality class. Weighted kappa statistics indicated that all three raters achieved good concordance (Figure 2B) in both the training (mean weighted-*κ* =0.64), internal testing (mean weighted-*κ* = 0.68), and external testing datasets (mean weighted-*κ* = 0.81). Additionally, polychoric correlations (Figure 2G-I), indicated very high correlation between raters in all datasets (training: mean *r* = 0.93; internal testing: mean *r* = 0.94; external testing datasets mean *r* =.94).

**Figure 2:**
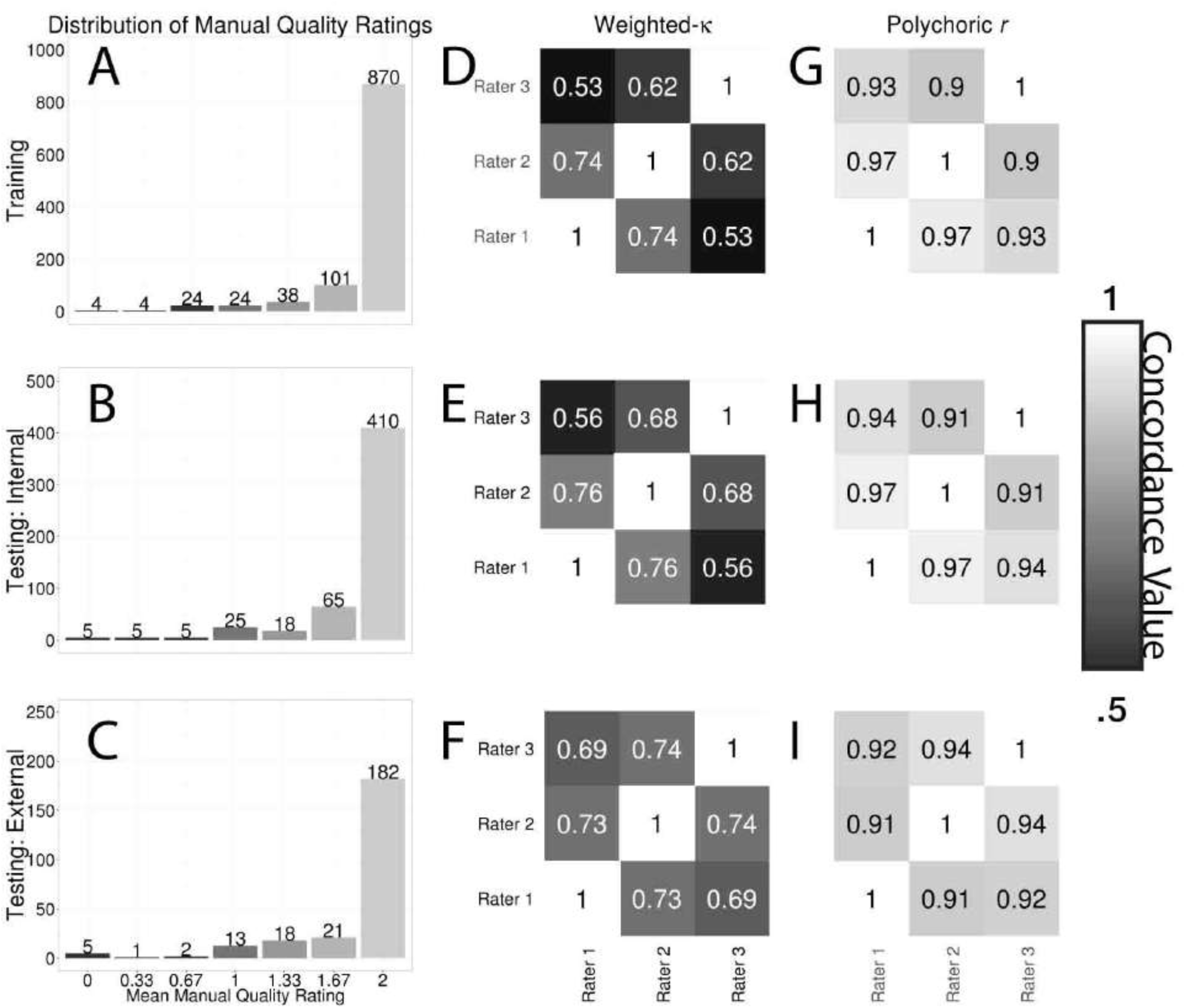
Results of manual ratings. ***A-C*:** Frequency of average manual rating for the training, internal testing, and external testing datasets. ***D-F*:** The pairwise weighted-κ between each rater in dataset was moderate and consistent across datasets. ***G-I*:** The pairwise polychoric correlation for each rater in all of the datasets was high.

### Manual quality ratings vary by age

While controlling for age, no sex differences were present in manual rating in any of the three datasets (training: *W* = 137700, p > 0.1, Figure 3A; internal testing: *W* = 36791, p > 0.1, Figure 3B; external testing: *W* = 6925, p > 0.1, Figure 3C). However, in both developmental samples from the PNC, younger age was associated with lower quality (training: ρ =  .14, p < 0.0001, Figure 3D; internal testing: ρ = .12, p < 0.01, Figure 3E). In contrast, among the older adults from the external testing dataset, greater age was associated with lower quality (ρ = -0.15, p < 0.05, Figure 3F).

**Figure 3:**
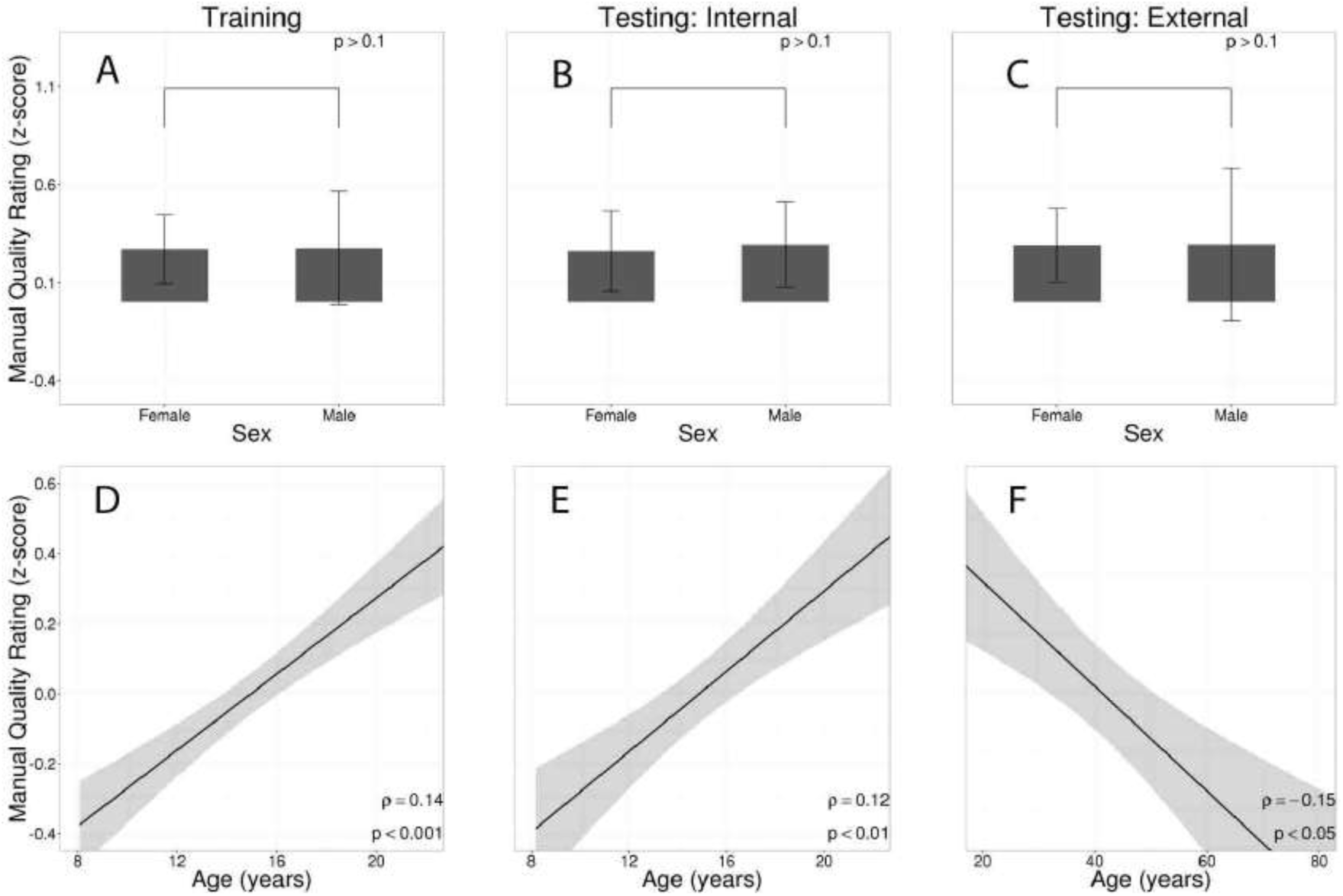
Manual quality rating varies by age but not by sex. No sex differences in median quality rating was observed in any of the three datasets (***A-C***); bars represent the median *z*-scored quality rating, error bars denote the inter-quartile range. Image quality improves with age during adolescence in both training (***D***) and internal testing samples (***E***) using PNC data, whereas data quality declines with aging over the adult lifespan in the external test dataset (***F***). In ***D-F,*** dark line represents a linear fit; shaded envelope represents 95% confidence intervals; reported significance values are calculated using partial Spearman’s correlations after regressing out gender trends.

### Quantitative measures of image quality align heterogeneously with manual rating

Next, we evaluated how quantitative measures of data quality related to the average quality rating across three raters. Putative quality measures displayed heterogeneous associations with manual quality ratings, both across measures and sometimes across datasets (Figure 4). The Euler number had the strongest association with manual rating across all three datasets. Furthermore, while the relationship was consistent across datasets for some measures (e.g., Euler number, Qi1), other measures were less consistent. For example, measures such as SNR, CNR, and FBER had only weak associations in the two PNC datasets, but had stronger associations in the external testing dataset that was acquired on a different scanner.

**Figure 4:**
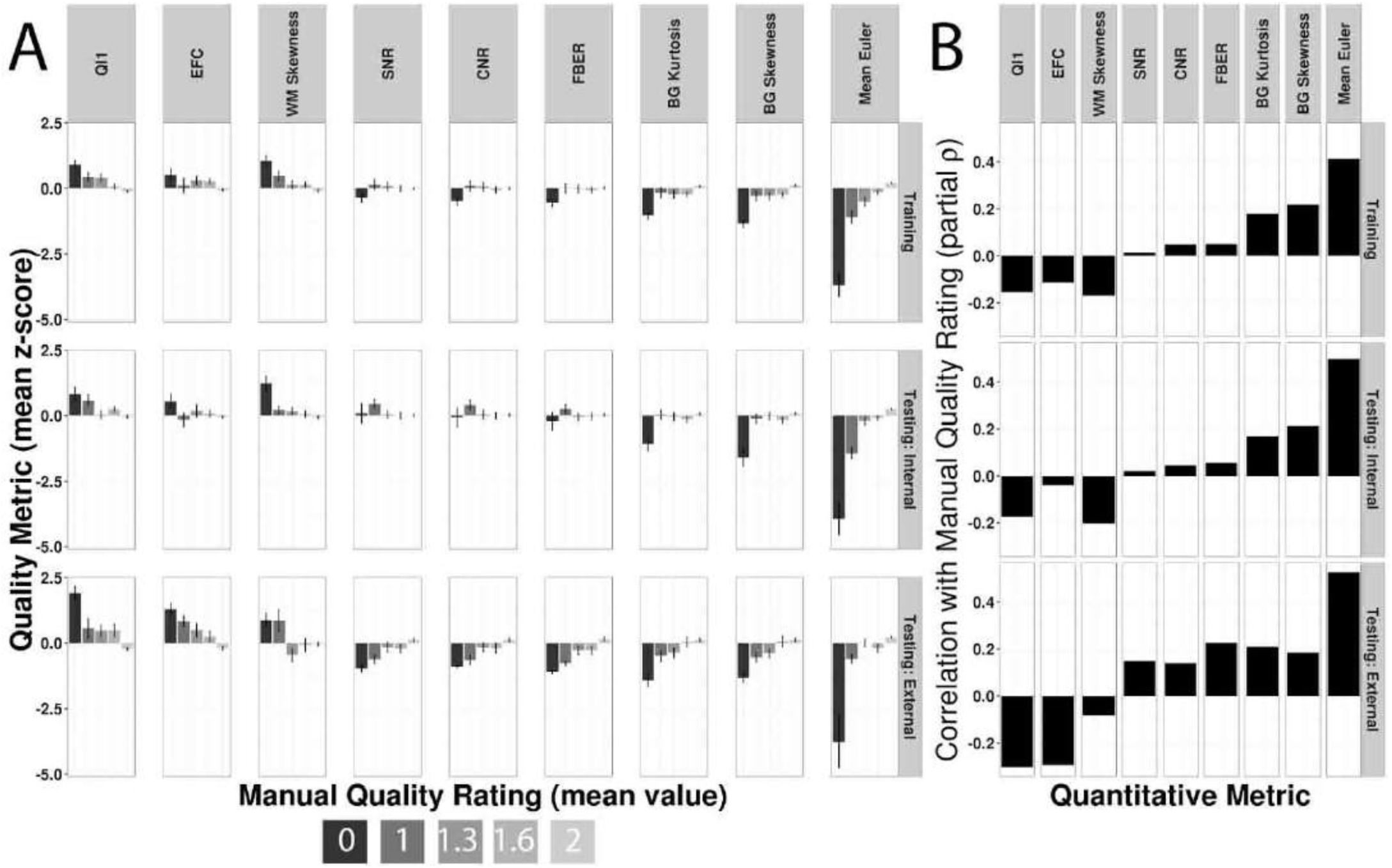
Quantitative metrics of image quality show heterogeneous alignment with manual ratings. ***A***: The standardized mean (+/- S.E.M.) for each quantitative metric is displayed by average manual rating class. ***B*:** Partial Spearman correlation coefficients between average manual quality rating and the T1 derived quantitative metrics; covariates included sex, age, and age squared. Across all datasets, Euler number showed the strongest association with manual quality ratings.

### Euler number successfully identifies unusable images

Next, we used the quantitative metrics to build an “inclusion” model that discriminated unusable images (rated “0”) from usable images (rated “1” or “2”). We began by measuring the classification capacity of each quantitative metric to identify a usable image (Figure 5A-C). Notably, the Euler number proved to be the most predictive feature across datasets (training: AUC = 0.99; internal testing: AUC = 0.98; external testing: AUC =0.99; Figure 5D-F). The Euler number value used for the classification threshold criteria were calculated using the training sample (accuracy = 0.94), and then applied to each test set. In the internal test set, accuracy remained quite high (accuracy = 0.92), but performance was somewhat lower in the external test set (accuracy = 0.76). Lower accuracy in the external test set was the result of very high sensitivity, but lower specificity (Table 3). As expected, the Euler number showed similar relationships to age and sex as the manual quality ratings (see Supplementary Figure 1).

**Figure 5:**
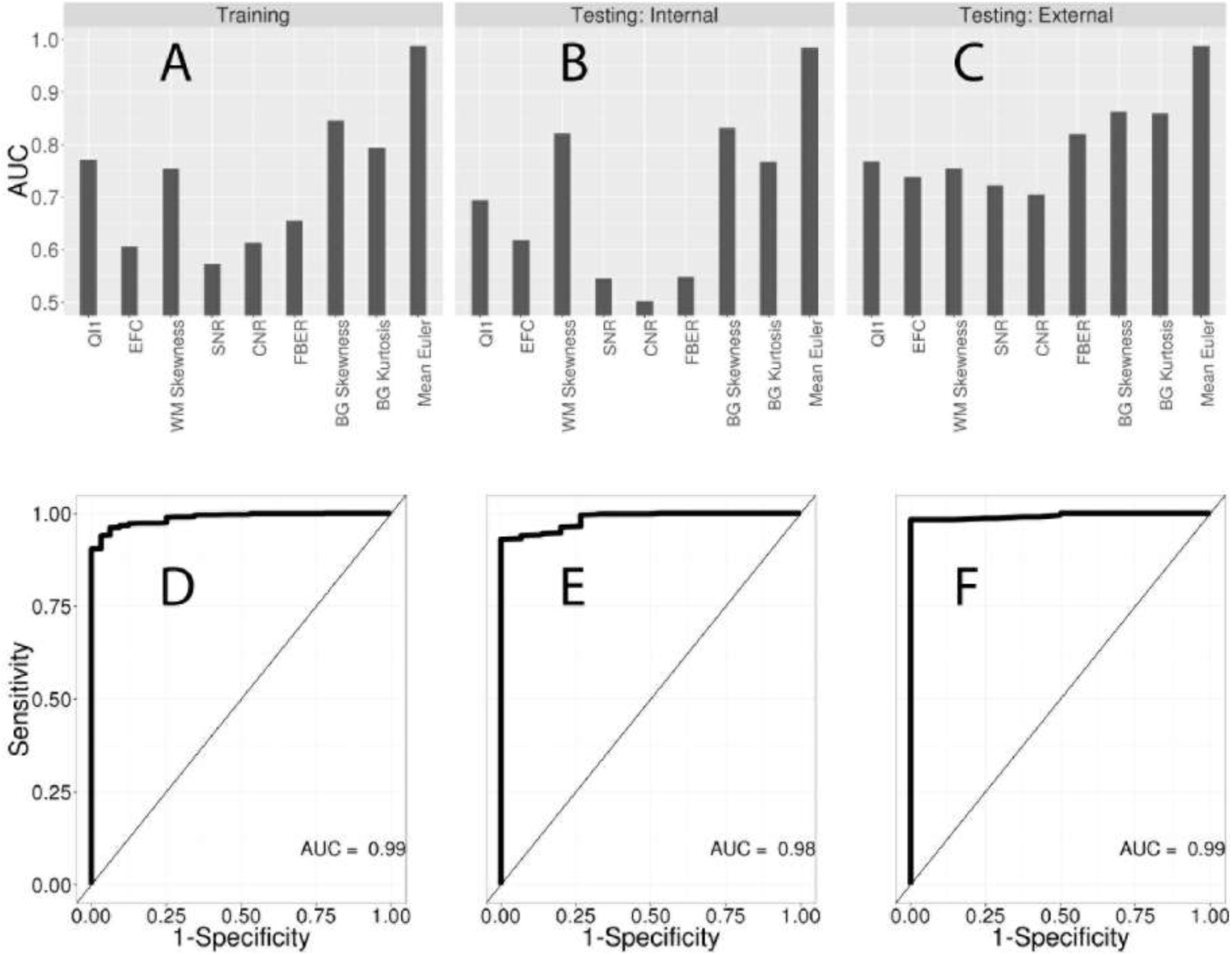
Inclusion model to identify unusable images. ***A-C*:** Logistic models in training (***A***), internal testing (***B***), and external testing (***C***) datasets were used to evaluate the ability of each quantitative measure of image quality to discriminate usable (rated 1-2) and unusable (rated 0) images. Area under the curve (AUC) was used to summarize model performance. In all datasets, the Euler number was the best-performing metric; adding additional metrics to the Euler number did not improve model performance. ***D-F***: Receiver Operator Characteristic (ROC) curves for the Euler number in each dataset.

**Table 3:** Inclusion model performance, using classification threshold criterion derived from training sample.

| Study | Threshold | Sensitivity | Specificity | Accuracy |
| --- | --- | --- | --- | --- |
| Training | -217 | 0.97 | 0.94 | 0.94 |
| Internal Testing | -217 | 1 | 0.93 | 0.92 |
| External Testing | -217 | 1 | 0.76 | 0.76 |

Notably, when the classification threshold criteria were allowed to vary by dataset, accuracy was quite high across all samples (range: 0.93-0.98; see Table 4). However, even when the threshold was varied by dataset, the inclusion model using the Euler number tended to be more sensitive than specific, with more false positives than false negatives. In this case, false positives were images flagged as unusable which were rated as usable by the manual raters. Post-hoc examination of these images revealed that, although they were not flagged as unusable by raters, these images did have a lower manual quality rating than those images which were marked as usable by both raters and the logistic model (training: n = 64, W = 35286, p < .1; internal testing: n = 40, W = 12452, p < 0.01; external testing: n = 57, W = 6952, p < 0.0001).

**Table 4:** Inclusion model performance, using classification threshold criterion calculated separately for each dataset.

| Study | Threshold | Sensitivity | Specificity | Accuracy |
| --- | --- | --- | --- | --- |
| Training | -217 | 0.97 | 0.94 | 0.94 |
| Internal Testing | -224.5 | 1 | 0.93 | 0.93 |
| External Testing | -380 | 1 | 0.98 | 0.98 |

### Limits of proxy measures from functional sequences

Based on prior reports that motion in functional sequences acquired in the same scanning session can provide a useful proxy of structural image quality, we next compared such proxy measures to those derived directly from the structural image. Specifically, we compared the Euler number to frame displacement from the four functional scans acquired as part of the PNC. As expected, motion within each sequence increased as the scanning session progressed (Figure 6A). Many participants did not complete all functional sequences, with more missing data for sequences acquired later in the session. Perhaps more importantly, attrition over the scanning session scaled directly with the data quality on the structural scan, such that those with lower structural image quality were less likely to have completed the subsequent functional sequences (Figure 6B). Furthermore, measures of motion during the functional sequences were less able to successfully identify unusable image compared to the Euler number (Figure 6C).

**Figure 6:**
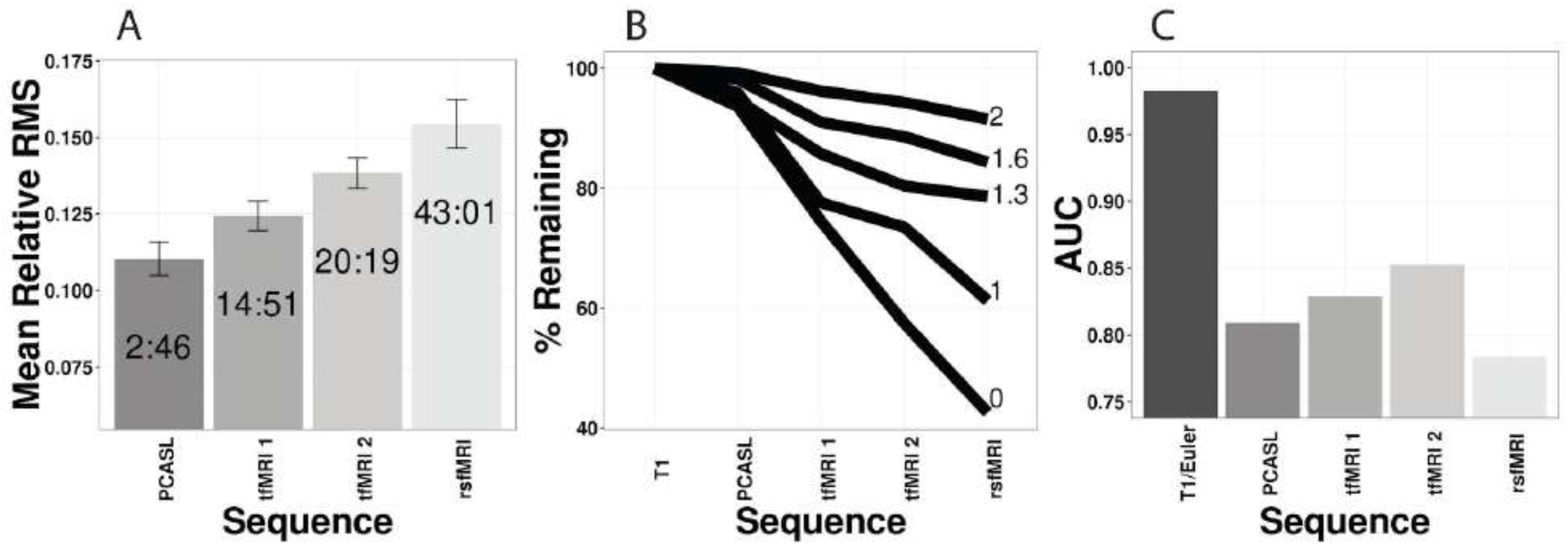
Limits of motion from functional scans as a proxy measure of T1 volume quality. ***A*:** Mean in-scanner motion during functional sequences acquired as part of the PNC increased over the course of the scanning session. Sequences are plotted in order of acquisition after the T1 scan; time from the T1 scan is reported in minutes: seconds within each bar. ***B*:** Individuals with lower-quality T1 images had differential attrition over the course of the of the scanning session. Thus, individuals with a lower-quality T1-images were less likely to complete the functional sequences which were subsequently acquired. Attrition scaled with quality of the T1 image. ***C*:** In participants for whom complete data was available (n=1275), motion estimated from the functional sequence did not perform as well as the Euler number in identifying unusable images (rated “0”).

### Quantitative estimates of data quality are related to cortical thickness

Having demonstrated that the Euler number can effectively identify unusable images (rated “0”), we next examined if this measure was related to cortical thickness in images that were considered usable (rated “1” or “2”). To do this, we conducted mass-univariate linear regression analyses evaluating the relationship between data quality (as summarized by the mean Euler number) with regional cortical thickness estimated using FreeSurfer. Across all three samples, highly consistent effects were observed. Overall, there was an FDR-corrected relationship with data quality in 53% of cortical regions (Figure 7A) in the training dataset, 44% of regions in the internal testing dataset(Figure 7B), and 39% of regions in the external testing dataset (Figure 7C). However, the directionality of this association was regionally heterogeneous. In regions including the dorsolateral prefrontal cortex, superior parietal cortex, and lateral temporal cortex, higher data quality was associated with thicker cortex. In contrast, in occipital and posterior cingulate cortex, higher data quality was associated with thinner cortex.

**Figure 7:**
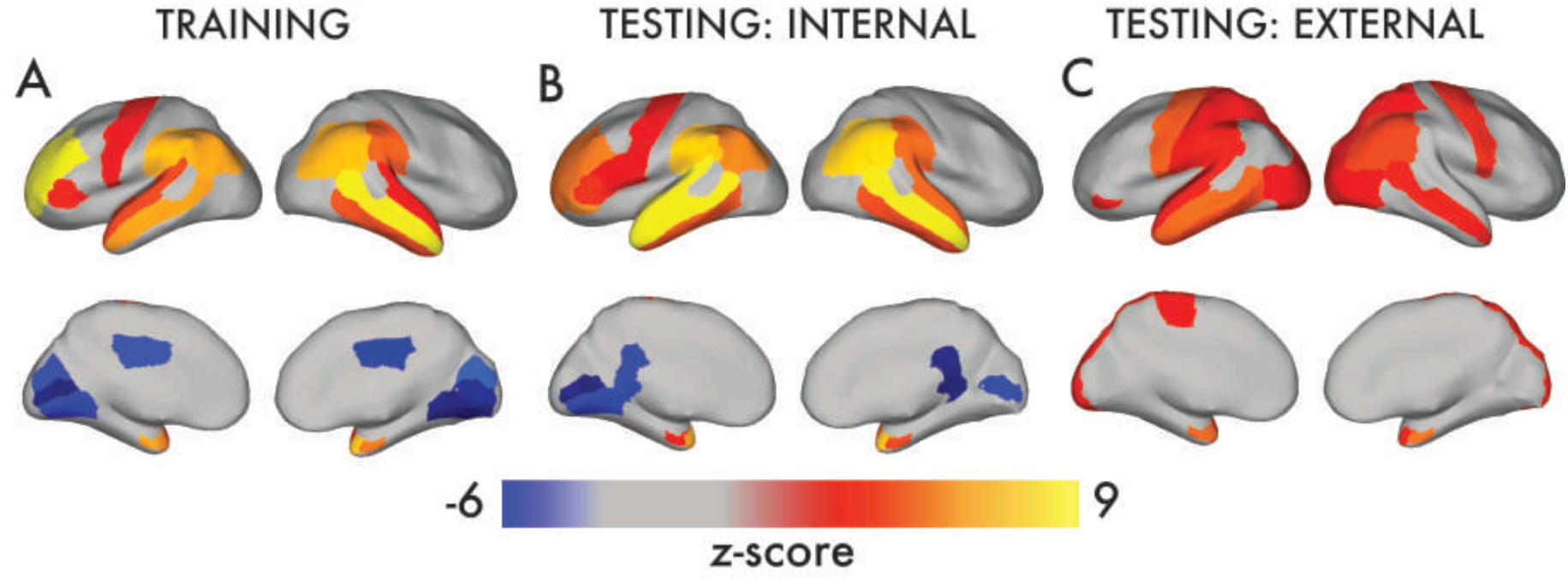
Quantitative measure of image quality is associated with cortical thickness. In usable images that were not excluded due to gross artifact, cortical thickness was significantly related to the Euler number in a regionally heterogeneous pattern. Higher data quality was associated with thicker cortex over much of the brain, but was conversely associated with thinner cortex in occipital and posterior cingulate cortex. This pattern was present across all datasets. Image displays *z*-scores from a mass-univariate linear regression, where regional cortical thickness was the outcome and Euler number was the predictor of interest; covariates included age, age squared, and sex. All results corrected for multiple comparisons using the False Discovery Rate (*q* < 0.05).

**Figure 8:**
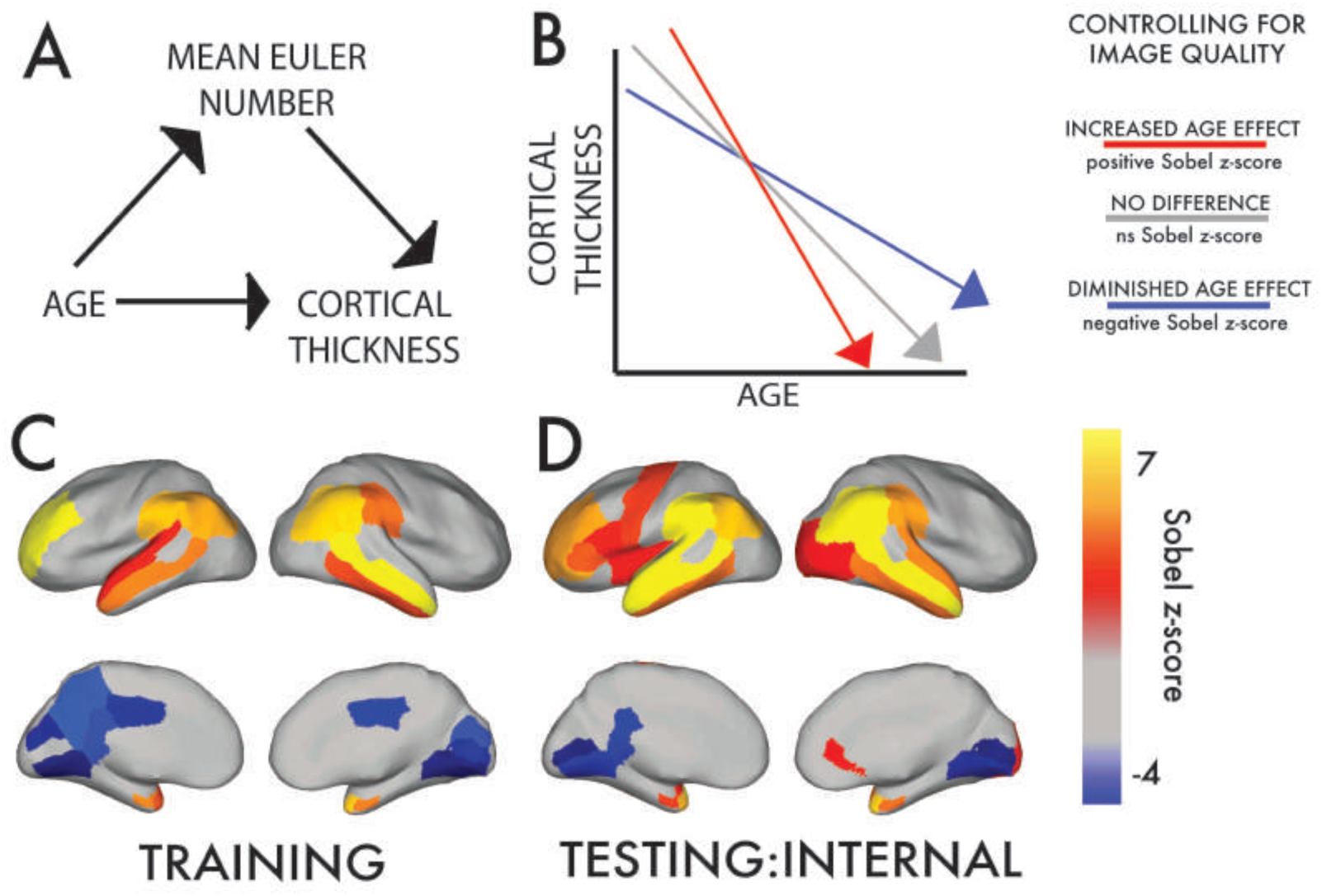
Data quality significantly mediates observed associations with age in youth. Having found that data quality is associated with both age and cortical thickness, we evaluated whether data quality might systematically bias inference regarding brain development. To do this, a mediation analysis was performed for each cortical region (***A***), where we evaluated if the Euler number mediated the apparent relationship between age and cortical thickness. At each region, Sobel *z*-scores were calculated as the test statistic for the mediation analysis. A positive Sobel’s value indicates that when controlling for data quality an increased effect of age was revealed; a negative Sobel’s value indicates that when controlling for data quality a diminished association with age was present (***B***). This procedure was applied to both the training (***C***) and internal test set (***D***) from the PNC, which revealed consistent mediation effects in both samples. Data quality significantly mediated the relationship between age and cortical thickness in a bidirectional, regionally heterogeneous manner. After controlling for data quality, the apparent age effect was increased in many regions (regions in warm colors), where higher data quality was associated with thicker cortex (see Figure 7). However, in a subset of regions including the occipital and posterior cingulate cortex, controlling for data quality resulted in a diminished association with age (cool colors). Multiple comparisons were accounted for using FDR (*q* <0.05).

### Data quality systematically biases associations with age in youth

The above results demonstrate that the Euler number aligns with manual ratings, is related to age, and is related to cortical thickness even among images considered usable. As a final step, we evaluated the degree to which data quality might bias inference regarding cross-sectional associations with age. Accordingly, we conducted mediation analyses to examine the degree to which data quality might mediate the relationship between age and brain structure (see schematics in **Figure 9A** & **B**). As expected given regionally heterogeneous effects of data quality on cortical thickness, data quality had a bidirectional impact on associations with age (**Figures 9C** & **D**). For most regions (shown in red), the relationship with data quality resulted in a masking of age effects, with observed associations with age becoming more significant when controlling for data quality. This reflects the fact that lower data quality makes the thick cortex of younger participants appear thinner, reducing estimates of thinning with age. In contrast, in several regions (shown in blue) including the posterior cingulate cortex and occipital cortex, data quality had the opposite effect, and inflated apparent age effects. Results were highly concordant in the training and internal testing datasets.

## DISCUSSION

In this paper, we demonstrate that a single quality measure derived from a T1-weighted volume – the Euler number – effectively recapitulates results from visual inspection with high accuracy. Furthermore, we demonstrate that image-based measures of data quality show differential relationships to several common measures of brain structure, and that data quality systematically biases associations between cortical thickness and age in youth.

### Manual raters can achieve a high level of concordance in a large-scale sample

It is increasingly recognized that data quality may be the primary confound in brain imaging studies of individual difference, lifespan development, or clinical populations (Ciric et al., 2017; Power et al., 2015). In-scanner motion is usually the single biggest determinant of data quality, especially in individuals who are young, elderly, or ill. While summary measures of motion can be easily derived from the realignment parameters of functional time series, motion cannot be easily estimated for most existing structural imaging data. A variety of motion-tracking and –correction systems have been developed (Zaitsev et al., 2015) However, such technologies have not been used for the vast majority of already-collected imaging data, which represents a huge societal investment. Due to the absence of a known ground truth, one of the first challenges for any study attempting to estimate image-derived measures of data quality for structural images is to create manual ratings, which are necessary to validate subsequent quantitative models. This problem is quite analogous to studies of psychiatric or neurologic illness, where several clinicians evaluate information from a patient and arrive at consensus diagnosis.

With limited training utilities available, we pursued an approach analogous to established procedures for training on clinical interviews and rating scales (Forbes et al., 2010; Kaufman et al., 1997). A panel of experts initially created a small set of anchors. Notably, while we originally piloted a rating system with 5 levels similar to that used in one recent study (Pardoe et al., 2016), we found that even highly trained experts could not reach a high level of concordance across 5 levels. Accordingly, we limited the quality rating to three levels, akin to previous efforts (Reuter et al., 2015; Savalia et al., 2017). Using these anchors, a larger training set of 100 images was then rated by two faculty experts. This set of 100 images was then used to train three experienced staff members to >85% accuracy. After this degree of reliability was established, the full set of images was evaluated. Following such training, concordance remained relatively good in both the training and testing samples. The pairwise correlation between raters was even higher, reflecting that when raters were not concordant it was usually due to a small but significant between-individual rater bias.

### The Euler number aligns with manual ratings and can identify unusable images

Having established a reliable set of manual ratings, we next derived quantitative measures of data quality using summary statistics from the structural image alone. Most of the measures we evaluated were produced using the Quality Assurance Pipeline (QAP)(Shehzad et al., 2015) included in the Configurable Pipeline for Analysis of Connectomes (C-PAC)(Sikka et al., 2013). In addition to this suite of measures, we also evaluated the Euler number, a measure of topological complexity of the cortical surface as reconstructed by FreeSurfer (Fischl, 2012). Using these measures, we examined the correspondence with the average quality rating across our three raters. Notably, the Euler number showed the highest correlation with the manual ratings across all three samples, suggesting it is a robust, dimensional measure of data quality.

In addition to being correlated with manual ratings, we also found that the Euler number was effective in identifying images that were so corrupted by artifact as to be unusable. This is a common step in sample construction in any imaging study. Notably, the Euler number had excellent performance across all three samples, with an AUC of 0.98-0.99. While AUC provides a good description of the overall predictive performance across all thresholds, a more stringent test of generalizability is whether a specific classification threshold from a model trained on one dataset can be applied to a different one. We found that a classification threshold which had excellent performance in the training data also performed quite well on the independent test set from the same study and scanner.

However, when this specific threshold was applied to an external test set, classification accuracy was substantially lower despite a near-perfect AUC (0.99). This reflects the fact that the specific classification threshold criteria from the training dataset of adolescents was not optimal for an adult sample acquired on a different scanner, and resulted in a very high sensitivity but lower-than-optimal specificity. However, when the threshold criteria were tailored to each dataset, performance was uniformly high. This suggests that the Euler number may be an effective measure of data quality across samples and scanners, but that the specific value used for flagging volumes for exclusion may need to be specified individually at each scanning site.

### Proxy measures of structural data quality from functional scans have important limits

One recently proposed approach is to use motion estimated from a functional time series acquired within the same session as a proxy of structural image quality. Several prior reports have shown that this is a fruitful approach (Alexander-Bloch et al., 2016; Pardoe et al., 2016; Savalia et al., 2017), demonstrating associations between this proxy measure of data quality and cortical thickness. One clear limitation of this approach is that it requires a functional scan to be acquired in the same scanning session. Furthermore, even when a functional scan is scheduled to be part of the imaging session, such data may be missing due to attrition. We demonstrated that motion increases over the course of the scanning session, and that participants with low-quality T1 volumes are more likely to be missing subsequent functional scans. Furthermore, our results show that frame displacement from functional scans are less able to identify unusable scans than the Euler number, which is calculated from the T1 volume itself.

### Measures of brain structure are differentially impacted by data quality

Previous work has shown that cortical thickness is systematically biased by in-scanner motion, whether quantified by manual rating (Pardoe et al., 2016; Savalia et al., 2017), motion estimated from functional sequences acquired in the same scanning session (Alexander-Bloch et al., 2016; Pardoe et al., 2016; Savalia et al., 2017), or volumetric navigators embedded in the T1 sequence (Reuter et al., 2015). Here, we demonstrate that an index of image quality derived directly from the structural image itself shows a similar relationship. Importantly, the association between data quality and cortical thickness had notable regional heterogeneity. In somatomotor, temporal, parietal, and many frontal regions, higher data quality was associated with greater thickness. However, in other regions including the visual cortex and posterior cingulate, higher data quality was associated with thinner estimated cortical thickness. These results are strikingly convergent with prior reports using other indices of data quality, which have demonstrated that while in general higher data quality is associated with thicker cortex, specific regions show the opposite effect (Alexander-Bloch et al., 2016; Pardoe et al., 2016; Reuter et al., 2015).

### Data quality biases estimates of structural brain development in youth

Accurate measurement of cortical thickness is critical to understanding typical and atypical trajectories of the developing brain. The extant literature indicates robust age-related cortical thinning in adolescence (Gennatas et al., 2017; Gogtay et al., 2004; Sowell et al., 2003, 2004; Tamnes et al., 2010). Moreover, there are regional-specific patterns of cortical maturation throughout development, with delayed maturation of higher-order association cortex (Giedd et al., 1999; Giedd, 2004; Gogtay et al., 2004; Shaw et al., 2008; Sowell et al., 2004; Tamnes et al., 2010). While most of these studies use validated methods to reduce in-scanner head motion during acquisition, few if any systematically evaluated or controlled for data quality. Importantly, several recent reports described significant relationships between age and in-scanner head motion in a variety of MRI protocols (Power et al., 2012; Roalf et al., 2016; Satterthwaite et al., 2016).

To determine if previously reported developmental trends are resilient to the impact of data quality, we performed region-wise mediation analyses. Notably, associations between cortical thickness and age were significantly mediated by data quality. This bias introduced by data quality was bidirectional and regionally heterogeneous. Several regions in frontal, temporal, parietal cortices showed more prominent developmental effects once T1 data quality was considered, suggesting that noise associated with data quality may partially mask associations with age. In contrast, regions such as the posterior cingulate, precuneus, and occipital cortex showed less prominent associations with age after controlling for data quality. These results emphasize that accurate delineation of cortical development is predicated upon data quality, which can both obscure important developmental effects in some regions and inflate effects in others. Notably, because data quality is likely to be collinear with other subject-level variables including cognitive performance (Siegel et al., 2017), symptom burden, and group status (Yendiki et al., 2014), this effect has the potential to similarly confound a wide variety of studies of brain structure.

### Limitations

Several important limitations of the current study should be noted. As discussed above, the Euler number provided an accurate image-based index of data quality across three datasets from two different scanners. However, the best exact classification threshold for accurate identification of unusable data did vary by scanner. Thus, one limitation of the current approach is that it is unlikely that a single Euler number exclusion threshold will apply to all studies. Second, in contrast to the measures provided by QAP, calculating the Euler number at present requires cortical surface reconstruction with FreeSurfer. This process is both time and computationally intensive, requiring 12-24 hours. This may limit the deployment of this index in certain settings. Moving forward, further investigation of other, simpler, registration-based methods may reveal that much of the same information can be gleaned from processes that are much less computationally demanding. However, given the widespread popularity of the FreeSurfer platform, it is also quite likely that many investigators have already calculated the Euler number for much of their data, allowing for immediate use in ongoing studies. Third, it is unknown at present how the test-retest reliability of automated measures of data quality (such as the Euler number) compare to manual ratings. However, in contrast to manual ratings, automated measures are 100% reproducible for a single image, and thus may also be more stable over time. Fourth, our quantitative quality metrics were selected according to their agreement with manual ratings. However, it should be acknowledged that manual ratings are not “ground truth” regarding image quality, and thus may be limited in their ability to inform and select quantitative quality measures. Fifth, due to our focus on cortical thickness, we did not evaluate the impact of data quality on sub-cortical or cerebellar regions. Finally, it should be noted that our use of the relatively coarse parcellation provided by the commonly-used Desikan-Killiany atlas precludes mapping the impact of data quality onto functional sub-systems (Gordon et al., 2016; Yeo et al., 2011).

### Conclusions

In this paper, we demonstrate that data quality can be estimated directly from structural images that lack volumetric navigators. Such image-based indices of data quality such as the Euler number can be used to exclude unusable images in a reproducible fashion. Furthermore, these continuous measures of image quality have the potential to be used as covariates in group-level analyses of structural imaging data. The ability to derive a measure of data quality directly from the structural image may obviate the need for use of proxy measures from functional sequences.

More broadly, the present data emphasize the degree to which data quality should be appreciated as an important confound in structural imaging studies. Investigators are encouraged to report measures of data quality for all structural imaging studies, especially those that evaluate individual or group differences. This is particularly relevant for studies where data quality is likely to be systematically related to the primary subject-level variable of interest, such as age, cognitive performance, clinical group status, or disease severity. We provide one such example, demonstrating that data quality can systematically bias associations between cortical thickness and age in youth. While it is now common practice to report summary measures of motion and image quality for fMRI research, it is less common for studies using T1-weighted imaging. The present results underscore a need for transparent reporting of such data. We urge investigators to report associations between data quality and both subject level variables of interest (e.g., age, group) as well as the primary imaging measure evaluated. Moving forward, quantitative estimates of motion during the T1 scan provided by motion-tracking and –correction technologies may obviate the need for post-hoc calculation of quality indices. However, we anticipate that the strategy outlined here may prove to be useful for the massive amount of structural imaging data that has already been collected at great effort and cost.

## ACKNOWLEDGEMENTS

We thank the acquisition and recruitment team, including Karthik Prabhakaran and Jeff Valdez. Thanks to Chad Jackson for data management and systems support. Supported by grants from the National Institute of Mental Health: R01MH107703 (TDS), R01MH112847 (TDS & RTS), R01MH107235 (RCG), R01MH112070 (CD), K01MH102609 (DRR), R01NS085211 (RTS), K01ES026840 (JES). Additional support was provided by the Dowshen Program for Neuroscience and the Penn/CHOP Lifespan Brain Institute. The PNC was funded through NIMH RC2 grants MH089983 and MH089924 (REG). Support for developing statistical analyses (RTS & TDS) was provided by a seed grant by the Center for Biomedical Computing and Image Analysis (CBICA) at Penn. Finally, we thank our anonymous reviewers for their valuable suggestions.

---

**Supplemental Figure 1:**
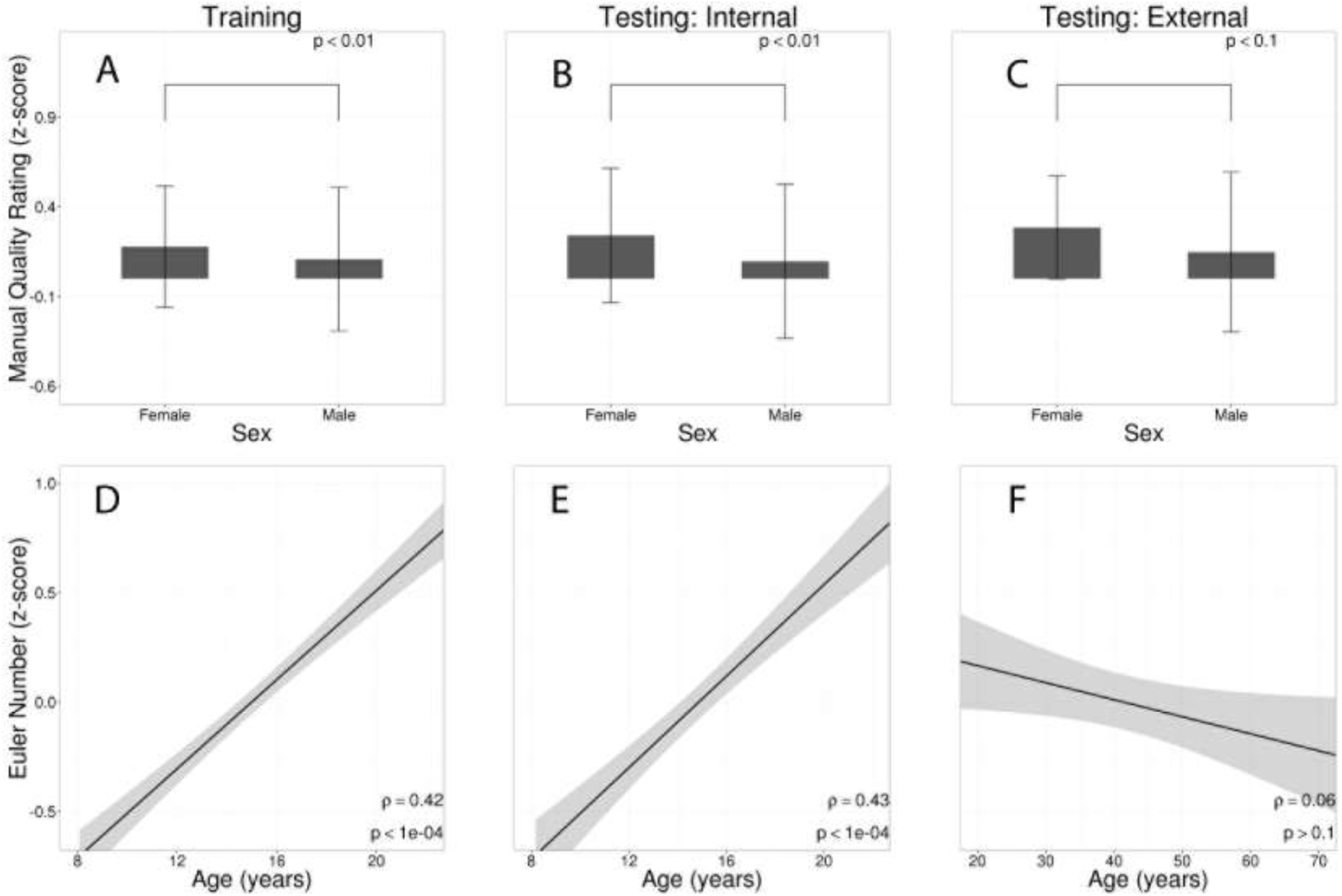
Mean Euler number varies by age and sex. Sex differences in medianquality rating was observed in all three datasets with females displaying higher Euler numbers (***A-C***); bars represent the median *z*-scored quality rating, error bars denote the inter-quartile range. Image quality improves with age during adolescence in both training (***D***) and internal testingsamples (***E***) using PNC data. However in the adult external testing sample a nonsignificant relationship between age and the Euler was observed although a negative trend was observed (***F***). In ***D-F,*** dark line represents a linear fit; shaded envelope represents 95% confidence intervals; reported significance values are calculated using partial Spearman’s correlations after regressingds. out gender trends.

## REFERENCES

Alexander-Bloch, A., Clasen, L., Stockman, M., Ronan, L., Lalonde, F., Giedd, J., Raznahan, A., 2016. Subtle in-scanner motion biases automated measurement of brain anatomy from in vivo MRI. Hum. Brain Mapp. 37, 2385–2397. doi:10.1002/hbm.23180

Atkinson, D., Hill, D.L., Stoyle, P.N., Summers, P.E., Keevil, S.F., 1997. Automatic correction of motion artifacts in magnetic resonance images using an entropy focus criterion. IEEE Trans. Med. Imaging 16, 903–910. doi:10.1109/42.650886

Avants, B.B., Kandel, B.M., Duda, J.T., Cook, P.A., Tustison, N.J., Shrinidhi, K., 2016. ANTsR: ANTs in R: quantification tools for biomedical images.

Bellon, E., Haacke, E., Coleman, P., Sacco, D., Steiger, D., Gangarosa, R., 1986. MR artifacts: a review. Am. J. Roentgenol. 147, 1271–1281. doi:10.2214/ajr.147.6.1271

Chen, J., Liu, J., Calhoun, V.D., Arias-Vasquez, A., Zwiers, M.P., Gupta, C.N., Franke, B., Turner, J.A., 2014. Exploration of scanning effects in multi-site structural MRI studies. J. Neurosci. Methods 230, 37–50. doi:10.1016/j.jneumeth.2014.04.023

Ciric, R., Wolf, D.H., Power, J.D., Roalf, D.R., Baum, G., Ruparel, K., Shinohara, R.T., Elliott, M.A., Eickhoff, S.B., Davatzikos, C., Gur, R.C., Gur, R.E., Bassett, D.S., Satterthwaite, T.D., 2017. Benchmarking of participant-level confound regression strategies for the control of motion artifact in studies of functional connectivity. NeuroImage. doi:10.1016/j.neuroimage.2017.03.020

Dale, A.M., Fischl, B., Sereno, M.I., 1999. Cortical surface-based analysis: I. Segmentation and surface reconstruction. Neuroimage 9, 179–194.

Dale, A.M., Fischl, B., Sereno, M.I., 1999. Cortical surface-based analysis. I. Segmentation and surface reconstruction. NeuroImage 9, 179–194. doi:10.1006/nimg.1998.0395

DeLong, E.R., DeLong, D.M., Clarke-Pearson, D.L., 1988. Comparing the areas under two or more correlated receiver operating characteristic curves: a nonparametric approach. Biometrics 44, 837–845.

Desikan, R.S., Ségonne, F., Fischl, B., Quinn, B.T., Dickerson, B.C., Blacker, D., Buckner, R.L., Dale, A.M., Maguire, R.P., Hyman, B.T., Albert, M.S., Killiany, R.J., 2006. An automated labeling system for subdividing the human cerebral cortex on MRI scans into gyral based regions of interest. NeuroImage 31, 968–980. doi:10.1016/j.neuroimage.2006.01.021

Fischl, B., 2012. FreeSurfer. NeuroImage 62, 774–781. doi:10.1016/j.neuroimage.2012.01.021

Fischl, B., Dale, A.M., 2000. Measuring the thickness of the human cerebral cortex from magnetic resonance images. Proc. Natl. Acad. Sci. U. S. A. 97, 11050–11055. doi:10.1073/pnas.200033797

Forbes, C., Blanchard, J.J., Bennett, M., Horan, W.P., Kring, A., Gur, R., 2010. Initial Development and Preliminary Validation of a New Negative Symptom Measure: The Clinical Assessment Interview for Negative Symptoms (CAINS). Schizophr. Res. 124, 36–42. doi:10.1016/j.schres.2010.08.039

Friston, K.J., Williams, S., Howard, R., Frackowiak, R.S.J., Turner, R., 1996. Movement-Related effects in fMRI time-series. Magn. Reson. Med. 35, 346–355. doi:10.1002/mrm.1910350312

Gennatas, E.D., Avants, B.B., Wolf, D.H., Satterthwaite, T.D., Ruparel, K., Ciric, R., Hakonarson, H., Gur, R.E., Gur, R.C., 2017. Age-Related Effects and Sex Differences in Gray Matter Density, Volume, Mass, and Cortical Thickness from Childhood to Young Adulthood. J. Neurosci. 37, 5065–5073. doi:10.1523/JNEUROSCI.3550-16.2017

Giedd, J.N., 2004. Structural magnetic resonance imaging of the adolescent brain. Ann. N. Y. Acad. Sci. 1021, 77–85. doi:10.1196/annals.1308.009

Giedd, J.N., Blumenthal, J., Jeffries, N.O., Castellanos, F.X., Liu, H., Zijdenbos, A., Paus, T., Evans, A.C., Rapoport, J.L., 1999. Brain development during childhood and adolescence: a longitudinal MRI study. Nat. Neurosci. 2, 861–863. doi:10.1038/13158

Gogtay, N., Giedd, J.N., Lusk, L., Hayashi, K.M., Greenstein, D., Vaituzis, A.C., Nugent, T.F., Herman, D.H., Clasen, L.S., Toga, A.W., Rapoport, J.L., Thompson, P.M., 2004. Dynamic mapping of human cortical development during childhood through early adulthood. Proc. Natl. Acad. Sci. U. S. A. 101, 8174–8179. doi:10.1073/pnas.0402680101

Gordon, E.M., Laumann, T.O., Adeyemo, B., Huckins, J.F., Kelley, W.M., Petersen, S.E., 2016. Generation and Evaluation of a Cortical Area Parcellation from Resting-State Correlations. Cereb. Cortex N. Y. N 1991 26, 288–303. doi:10.1093/cercor/bhu239

Harvard Center for Brain Science, 2014. Quality Control.

Jenkinson, M., Bannister, P., Brady, M., Smith, S., 2002. Improved optimization for the robust and accurate linear registration and motion correction of brain images. NeuroImage 17, 825–841.

Joanes, D.N., Gill, C.A., 1998. Comparing measures of sample skewness and kurtosis. J. R. Stat. Soc. Ser. Stat. 47, 183–189. doi:10.1111/1467-9884.00122

Kaufman, J., Birmaher, B., Brent, D., Rao, U., Flynn, C., Moreci, P., Williamson, D., Ryan, N., 1997. Schedule for Affective Disorders and Schizophrenia for School-Age Children-Present and Lifetime Version (K-SADS-PL): initial reliability and validity data. J. Am. Acad. Child Adolesc. Psychiatry 36, 980–988. doi:10.1097/00004583-199707000-00021

Kuhn, M., Weston, S., Williams, A., Keefer, C., Engelhardt, A., Cooper, T., Mayer, Z., Kenkel, B., Team, the R.C., Benesty, M., Lescarbeau, R., Ziem, A., Scrucca, L., Tang, Y., Candan, C., Hunt, and T., 2016. caret: Classification and Regression Training.

Magnotta, V.A., Friedman, L., 2006. Measurement of Signal-to-Noise and Contrast-to-Noise in the fBIRN Multicenter Imaging Study. J. Digit. Imaging 19, 140–147. doi:10.1007/s10278-006-0264-x

Marcus, D.S., Harms, M.P., Snyder, A.Z., Jenkinson, M., Wilson, J.A., Glasser, M.F., Barch, D.M., Archie, K.A., Burgess, G.C., Ramaratnam, M., Hodge, M., Horton, W., Herrick, R., Olsen, T., McKay, M., House, M., Hileman, M., Reid, E., Harwell, J., Coalson, T., Schindler, J., Elam, J.S., Curtiss, S.W., Van Essen, D.C., 2013. Human Connectome Project Informatics: quality control, database services, and data visualization. NeuroImage 80. doi:10.1016/j.neuroimage.2013.05.077

Mortamet, B., Bernstein, M.A., Jack, C.R., Gunter, J.L., Ward, C., Britson, P.J., Meuli, R., Thiran, J.-P., Krueger, G., Alzheimer’s Disease Neuroimaging Initiative, 2009. Automatic quality assessment in structural brain magnetic resonance imaging. Magn. Reson. Med. 62, 365–372. doi:10.1002/mrm.21992

Pardoe, H.R., Kucharsky Hiess, R., Kuzniecky, R., 2016. Motion and morphometry in clinical and nonclinical populations. NeuroImage 135, 177–185. doi:10.1016/j.neuroimage.2016.05.005

Power, J.D., Barnes, K.A., Snyder, A.Z., Schlaggar, B.L., Petersen, S.E., 2012. Spurious but systematic correlations in functional connectivity MRI networks arise from subject motion. Neuroimage 59, 2142–2154. doi:10.1016/j.neuroimage.2011.10.018

Power, J.D., Schlaggar, B.L., Petersen, S.E., 2015. Recent progress and outstanding issues in motion correction in resting state fMRI. NeuroImage 105, 536–551. doi:10.1016/j.neuroimage.2014.10.044

Reuter, M., Tisdall, M.D., Qureshi, A., Buckner, R.L., van der Kouwe, A.J.W., Fischl, B., 2015. Head motion during MRI acquisition reduces gray matter volume and thickness estimates. NeuroImage 107, 107–115. doi:10.1016/j.neuroimage.2014.12.006

Revelle, W., 2017. psych: Procedures for Psychological, Psychometric, and Personality Research.

Roalf, D.R., Quarmley, M., Elliott, M.A., Satterthwaite, T.D., Vandekar, S.N., Ruparel, K., Gennatas, E.D., Calkins, M.E., Moore, T.M., Hopson, R., Prabhakaran, K., Jackson, C.T., Verma, R., Hakonarson, H., Gur, R.C., Gur, R.E., 2016. The impact of quality assurance assessment on diffusion tensor imaging outcomes in a large-scale population-based cohort. NeuroImage 125, 903–919. doi:10.1016/j.neuroimage.2015.10.068

Roalf, D.R., Vandekar, S.N., Almasy, L., Ruparel, K., Satterthwaite, T.D., Elliott, M.A., Podell, J., Gallagher, S., Jackson, C.T., Prasad, K., Wood, J., Pogue-Geile, M.F., Nimgaonkar, V.L., Gur, R.C., Gur, R.E., 2015. Heritability of Subcortical and Limbic Brain Volume and Shape in Multiplex-Multigenerational Families with Schizophrenia. Biol. Psychiatry 77, 137–146. doi:10.1016/j.biopsych.2014.05.009

Satterthwaite, T.D., Connolly, J.J., Ruparel, K., Calkins, M.E., Jackson, C., Elliott, M.A., Roalf, D.R., Hopson, R., Prabhakaran, K., Behr, M., Qiu, H., Mentch, F.D., Chiavacci, R., Sleiman, P.M.A., Gur, R.C., Hakonarson, H., Gur, R.E., 2016. The Philadelphia Neurodevelopmental Cohort: A publicly available resource for the study of normal and abnormal brain development in youth. NeuroImage, Sharing the wealth: Brain Imaging Repositories in 2015 124, Part B, 1115–1119. doi:10.1016/j.neuroimage.2015.03.056

Satterthwaite, T.D., Elliott, M.A., Ruparel, K., Loughead, J., Prabhakaran, K., Calkins, M.E., Hopson, R., Jackson, C., Keefe, J., Riley, M., Mentch, F.D., Sleiman, P., Verma, R., Davatzikos, C., Hakonarson, H., Gur, R.C., Gur, R.E., 2014. Neuroimaging of the Philadelphia neurodevelopmental cohort. NeuroImage 86, 544–553. doi:10.1016/j.neuroimage.2013.07.064

Satterthwaite, T.D., Wolf, D.H., Loughead, J., Ruparel, K., Elliott, M.A., Hakon, H., Gur, R.C., Gur, R.E., 2012. Impact of In-Scanner Head Motion on Multiple Measures of Functional Connectivity: Relevance for Studies of Neurodevelopment in Youth. NeuroImage 60, 623–632. doi:10.1016/j.neuroimage.2011.12.063

Savalia, N.K., Agres, P.F., Chan, M.Y., Feczko, E.J., Kennedy, K.M., Wig, G.S., 2017. Motion-related artifacts in structural brain images revealed with independent estimates of in-scanner head motion. Hum. Brain Mapp. 38, 472–492. doi:10.1002/hbm.23397

Shaw, P., Kabani, N.J., Lerch, J.P., Eckstrand, K., Lenroot, R., Gogtay, N., Greenstein, D., Clasen, L., Evans, A., Rapoport, J.L., Giedd, J.N., Wise, S.P., 2008. Neurodevelopmental trajectories of the human cerebral cortex. J. Neurosci. Off. J. Soc. Neurosci. 28, 3586–3594. doi:10.1523/JNEUROSCI.5309-07.2008

Shehzad, Z., Giavasis, S., Li, Q., Benhajali, Y., Yan, C., Yang, Z., Milham, M., Bellec, P., Craddock, C., n.d. The Preprocessed Connectomes Project Quality Assessment Protocol - a resource for measuring the quality of MRI data. Front. Neurosci. doi:10.3389/conf.fnins.2015.91.00047

Siegel, J.S., Mitra, A., Laumann, T.O., Seitzman, B.A., Raichle, M., Corbetta, M., Snyder, A.Z., 2017. Data Quality Influences Observed Links Between Functional Connectivity and Behavior. Cereb. Cortex N. Y. N 1991 27, 4492–4502. doi:10.1093/cercor/bhw253

Sikka, S., Cheung, B., Khanuja, R., Ghosh, S., Yan, C., Li, Q., Vogelstein, J., Burns, R., Colcombe, S., Craddock, C., Mennes, M., Kelly, C., Dimartino, A., Castellanos, F., Milham, M., n.d. Towards Automated Analysis of Connectomes: The Configurable Pipeline for the Analysis of Connectomes (C-PAC). Front. Neuroinformatics. doi:10.3389/conf.fninf.2014.08.00117

Smith, T.B., Nayak, K.S., 2010. MRI artifacts and correction strategies. Imaging Med. 2, 445–457. doi:10.2217/iim.10.33

Sobel, M.E., 1982. Asymptotic Confidence Intervals for Indirect Effects in Structural Equation Models. Sociol. Methodol. 13, 290–312. doi:10.2307/270723

Sowell, E.R., Peterson, B.S., Thompson, P.M., Welcome, S.E., Henkenius, A.L., Toga, A.W., 2003. Mapping cortical change across the human life span. Nat. Neurosci. 6, 309–315. doi:10.1038/nn1008

Sowell, E.R., Thompson, P.M., Leonard, C.M., Welcome, S.E., Kan, E., Toga, A.W., 2004. Longitudinal mapping of cortical thickness and brain growth in normal children. J. Neurosci. Off. J. Soc. Neurosci. 24, 8223–8231. doi:10.1523/JNEUROSCI.1798-04.2004

Styner, M.A., Charles, H.C., Park, J., Gerig, G., 2002. Multisite validation of image analysis methods: assessing intra- and intersite variability. pp. 278–286. doi:10.1117/12.467167

Tamnes, C.K., Ostby, Y., Fjell, A.M., Westlye, L.T., Due-Tønnessen, P., Walhovd, K.B., 2010. Brain maturation in adolescence and young adulthood: regional age-related changes in cortical thickness and white matter volume and microstructure. Cereb. Cortex N. Y. N 1991 20, 534–548. doi:10.1093/cercor/bhp118

Tisdall, M.D., Hess, A.T., Reuter, M., Meintjes, E.M., Fischl, B., van der Kouwe, A.J.W., 2012. Volumetric navigators for prospective motion correction and selective reacquisition in neuroanatomical MRI. Magn. Reson. Med. 68, 389–399. doi:10.1002/mrm.23228

Tisdall, M.D., Reuter, M., Qureshi, A., Buckner, R.L., Fischl, B., van der Kouwe, A.J.W., 2016. Prospective motion correction with volumetric navigators (vNavs) reduces the bias and variance in brain morphometry induced by subject motion. NeuroImage 127, 11–22. doi:10.1016/j.neuroimage.2015.11.054

Van Dijk, K.R.A., Sabuncu, M.R., Buckner, R.L., 2012. The influence of head motion on intrinsic functional connectivity MRI. NeuroImage 59, 431–438. doi:10.1016/j.neuroimage.2011.07.044

Wang, B., 2015. bda: Density Estimation for Grouped Data.

Woodard, J.P., Carley-Spencer, M.P., 2006. No-Reference image quality metrics for structural MRI. Neuroinformatics 4, 243–262. doi:10.1385/NI:4:3:243

Xu, T., Yang, Z., Jiang, L., Xing, X.-X., Zuo, X.-N., 2015. A Connectome Computation System for discovery science of brain. Sci. Bull. 60, 86–95. doi:10.1007/s11434-014-0698-3

Yan, C.-G., Cheung, B., Kelly, C., Colcombe, S., Craddock, R.C., Di Martino, A., Li, Q., Zuo, X.-N., Castellanos, F.X., Milham, M.P., 2013. A comprehensive assessment of regional variation in the impact of head micromovements on functional connectomics. NeuroImage 76, 183–201. doi:10.1016/j.neuroimage.2013.03.004

Yan, C.-G., Wang, X.-D., Zuo, X.-N., Zang, Y.-F., 2016. DPABI: Data Processing & Analysis for (Resting-State) Brain Imaging. Neuroinformatics 14, 339–351. doi:10.1007/s12021-016-9299-4

Yendiki, A., Koldewyn, K., Kakunoori, S., Kanwisher, N., Fischl, B., 2014. Spurious group differences due to head motion in a diffusion MRI study. NeuroImage 88, 79–90. doi:10.1016/j.neuroimage.2013.11.027

Yeo, B.T.T., Krienen, F.M., Sepulcre, J., Sabuncu, M.R., Lashkari, D., Hollinshead, M., Roffman, J.L., Smoller, J.W., Zöllei, L., Polimeni, J.R., Fischl, B., Liu, H., Buckner, R.L., 2011. The organization of the human cerebral cortex estimated by intrinsic functional connectivity. J. Neurophysiol. 106, 1125–1165. doi:10.1152/jn.00338.2011

Zaitsev, M., Maclaren, J., Herbst, M., 2015. Motion artifacts in MRI: A complex problem with many partial solutions. J. Magn. Reson. Imaging 42, 887–901. doi:10.1002/jmri.24850

Zhang, Y., Brady, M., Smith, S., 2001. Segmentation of brain MR images through a hidden Markov random field model and the expectation-maximization algorithm. IEEE Trans. Med. Imaging 20, 45–57. doi:10.1109/42.906424

Zuo, X.-N., He, Y., Betzel, R.F., Colcombe, S., Sporns, O., Milham, M.P., 2017. Human Connectomics across the Life Span. Trends Cogn. Sci. 21, 32–45. doi:10.1016/j.tics.2016.10.005

